# Episodic Control as Meta-Reinforcement Learning

**DOI:** 10.1101/360537

**Authors:** S Ritter, JX Wang, Z Kurth-Nelson, M Botvinick

**Affiliations:** DeepMind, London, UK; Princeton Neuroscience Institute, Princeton, NJ; MPS-UCL Centre for Computational Psychiatry, London, UK; Gatsby Computational Neuroscience Unit, UCL, London, UK

**Keywords:** Reinforcement learning, model-based, deep learning, meta-learning, episodic memory

## Abstract

Recent research has placed episodic reinforcement learning (RL) alongside model-free and model-based RL on the list of processes centrally involved in human reward-based learning. In the present work, we extend the unified account of model-free and model-based RL developed by Wang et al. (2018) to further integrate episodic learning. In this account, a generic model-free “meta-learner” learns to deploy and coordinate among all of these learning algorithms. The meta-learner learns through brief encounters with many novel tasks, so that it *learns to learn* about new tasks. We show that when equipped with an episodic memory system inspired by theories of reinstatement and gating, the meta-learner learns to use the episodic and model-based learning algorithms observed in humans in a task designed to dissociate among the influences of various learning strategies. We discuss implications and predictions of the model.

## Introduction

Nearly every decision an intelligent organism makes is informed by its memory of the results of its past decisions. To be successful, agents must distill the results of past decisions into memories, then make use of those memories to make better decisions in the future. Accordingly, much effort has been directed toward understanding (1) what humans and animals store after a sequence of actions and rewards, and (2) how they use that stored information to appraise the value of future actions.

Model-free and model-based reinforcement learning (RL; Daw, Gershman, Seymour, Dayan, & Dolan, 2011; Sutton & Barto, 1998) offer distinct solutions to these two problems. Model-free RL stores statistics about the relationship between states, actions and rewards, and appraises actions by calculating how frequently they led to reward. Meanwhile, model-based RL stores estimated state-state transition probabilities, and appraises actions by using this model to simulate sequences of states to predict future reward. Signatures of both model-free and model-based learning appear in behavior and in the brain (e.g. Daw et al., 2011), and a venerable tradition holds that they are implemented by dissociable neural systems (for review see Dolan & Dayan, 2013).

However, the recent theory of meta-reinforcement learning (meta-RL) proposed that model-free learning, model-based learning, and their sometimes complex interaction could all be explained by a simple unified mechanism (Wang et al., 2018). In meta-RL, a recurrent neural network (RNN) receives a reward signal as part of its input and is trained by model-free learning on a series of interrelated tasks to rapidly learn from this signal. Through this training, the RNN learns to distill the history of observations, actions, and rewards into its hidden state (a form of *working memory*) and to use this summary to select rewarding actions. In essence, this end result constitutes a learned reinforcement learning algorithm that operates in the RNN’s activation dynamics. Critically, Wang et al. (2017) showed that this meta-learned RL algorithm can be model-based, even though it was acquired through model-free learning.

While meta-RL provides a full account of incremental learning as it is carried out in working memory, it does not account for the episodic learning processes to which attention has recently been called (Gershman & Daw, 2017). In addition to learning by incrementally storing recent sequences of behavior in working memory, humans appear to learn by storing summaries of individual episodes for long periods of time, then retrieving them when similar contexts are encountered. For example, cues triggering episodic memory retrieval impact reward-based learning, both for good and for ill (Bornstein, Khaw, Shohamy, & Daw, 2017; Vikbladh, Shohamy, & Daw, 2017; Bornstein & Norman, 2017), and distinctive aspects of episodic memory function contribute to decision-making behavior (Wimmer, Braun, Daw, & Shohamy, 2014; Wimmer & Buechel, 2016; Duncan & Shohamy, 2016). Such observations, along with some fundamental computational insights (Lengyel & Dayan, 2007), have recently landed episodic learning a spot beside incremental model-free and model-based reinforcement learning on the list of processes centrally involved in decision making (Gershman & Daw, 2017).

In the present work, we develop a natural extension to meta-RL that enables it to integrate episodic learning. The resulting theory, based on an algorithm introduced to the machine learning literature by Ritter et al. (2018), explains how incremental and episodic learning, as well as the coordination between them can be meta-learned through purely model-free RL. The episodic meta-RL theory proposes the following:

1. Meta-RL’s working memory is supplemented by an episodic memory which stores working memory states.
2. Each state is paired with a perceptual context embedding that is later used to retrieve the working memory state when similar perceptual contexts are encountered.
3. The retrieved states are then gated into the working memory using a parameterized function, whose parameters are optimized toward the same model-free objective that trains the working memory dynamics.

This proposal is inspired in part by evidence that episodic memory retrieval in humans operates through reinstatement, triggering patterns of neural activity related to those that were induced by the original encoding of the relevant episode (e.g., Xiao et al., 2017), and evidence that reinstatement occurs not only in perceptual systems, but also recreates patterns of activity in neural circuits supporting working memory (Hoskin, Bornstein, Norman, & Cohen, 2017; Cohen & O’Reilly, 1996). Our implementation of this proposal draws additional inspiration from recent work on differentiable memory systems (e.g., Graves et al., 2016), especially that of Pritzel et al. (2017), which makes use of context-based retrieval for RL.

To empirically test this model, in this work we compared its behavior to that of humans observed by Vikbladh et al. (2017) in a task designed to dissociate the effects of multiple types of incremental and episodic learning. Vikbladh and colleagues found evidence of the use of a model-based form of episodic memory, whereby traces of specific episodes are retrieved from long-term memory based on visual similarity, then used along with knowledge of the transition structure of the environment to select actions. This episodic model-based learning was present in conjunction with incremental model-free and incremental model-based learning. In the following sections, we describe the task in detail and demonstrate that meta-RL with episodic memory reproduces this model-based episodic learning and its coordination with incremental model-based learning. To conclude, we consider directions for future work, including testable predictions of the theory.

## Task

The task we study is a version of the two-step task (Daw et al., 2011) augmented with episodic cues to previous trials. The task structure, which was inspired by Vikbladh et al. (2017), is diagrammed in Figure 1. Each trial consisted of two stages. On the first stage, state *s*_0_, the agent was either presented with the “no-cue” stimulus (a vector of all −1’s) on uncued trials, or with a binary vector associated with a previously seen second-stage context on cued trials (see Figure 1). In response, the agent chose either *a*_1_ or *a*_2_ and transitioned into one of two second-stage states *s*_1_ or *s*_2_ with probabilities *p*(*s*_1_|*a*_1_) = *p*(*s*_2_|*a*_2_) = 0.9 (common transition) and *p*(*s*_1_|*a*_2_) = *p*(*s*_2_|*a*_1_) = 0.1 (uncommon transition). These transition probabilities *p* were fixed across episodes. On the second stage, the agent was presented with a stimulus representing the context of that second-stage state, followed by a final step in which it was shown the reward outcome. On uncued trials, the states *s*_1_ and *s*_2_ yielded Bernoulli probabilistic rewards of 0 or 1 according to [*r_a_, r_b_*] = [0.9,0.1] or [0.1,0.9], with the reward contingencies having a 10% chance of switching at the beginning of each trial.

**Figure 1:**
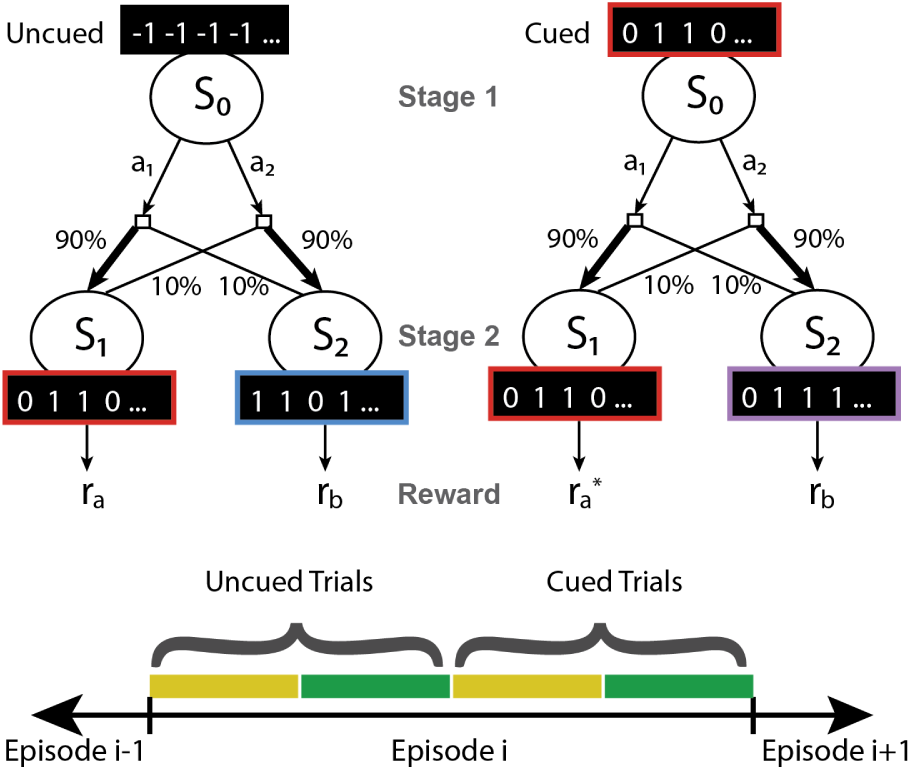
Contextual two-step task modeled after Daw et al. (2011) and Vikbladh et al. (2017). (Top) Two trial types are shown: uncued and cued. All trials start in state *s*_0_ at the first stage, at which point agents are presented with either a “no-cue” stimulus or are cued with a second-stage stimulus seen on a previous trial. Transition probabilities after taking actions *a*_1_ or *a*_2_ are depicted in the graph. On uncued trials, *s*_1_ and *s*_2_ result in Bernoulli rewards with probabilities *r_a_* and *r_b_*. On cued trials, transitioning into the same state as in the trial k during which the cue was first presented results in receiving the same reward received on trial k, 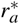. (Bottom) Trials within an episode are split into 4 blocks, with block 3 consisting of cued trials which are cued with stimuli from block 1, and block 4 cued from block 2.

On cued trials, if the agent transitioned into the same state as on the trial k during which the cue was first presented, the agent was given the exact same reward as on trial k. If the agent transitioned into the other state, the reward was determined as on uncued trials. This is the critical feature of the task for assessing episodic memory use; if the agent could remember the state entered and reward achieved on the cue-associated trial, it could act to access or avoid that state, depending on whether the past trial was rewarded.

The first half of every episode (50 trials) consisted entirely of uncued trials, and the second half consisted entirely of cued trials, with trials 51-75 being cued with stimuli from trials 1-25 and trials 76-100 cued with stimuli from trials 26-50, randomly sampled without replacement. This was done to reduce autocorrelation in the reward probabilities by enforcing a minimum of 25 trials between seeing the stimulus on the second stage and being cued with it on the first stage. The agent was trained for 10,000 episodes of 100 trials each, and evaluated with weights fixed on 500 further episodes.

### Learning Algorithms

The two-step task with episodic cueing is designed to dissociate among the influences of four different learning strategies on choice. First, the incremental model-free strategy prescribes taking the same action that was taken on the last trial if it was rewarded, and taking the opposite action if it was not rewarded, regardless of whether the transition on the previous trial was common or uncommon. In contrast, the incremental model-based strategy prescribes taking the same action only if the previous trial was rewarded *and* the previous transition was common. If the previous transition was uncommon and the trial was rewarded, the agent will take the opposite action. Episodic model-free and model-based strategies operate like their incremental counterparts, but with respect to the trial associated with the cue rather than the immediately previous trial.

## Model

In episodic meta-RL (EMRL; Figure 2; Ritter et al., 2018), vectors represent working memory states, and a recurrent neural network, specifically a long short-term memory network (LSTM; Hochreiter & Schmidhuber, 1997), updates the working memory state at each time step and uses it to select actions. To implement context-based reinstatement of the LSTM’s activations, EMRL augments this working memory with an episodic memory containing working memory states. Based on Pritzel et al.’s (2017) differentiable neural dictionary (see also Blundell et al., 2016), this episodic memory stores a visual representation of the context along with each item, which is used at retrieval time to find working memory states stored in contexts similar to the retrieval context. In our experiments, the EMRL agent writes to the memory at the end of the each trial. The agent reads from the memory on every time step by searching for the perceptual representation in the array with the smallest cosine distances from the representation of the current state, then retrieving the associated working memory state.

**Figure 2:**
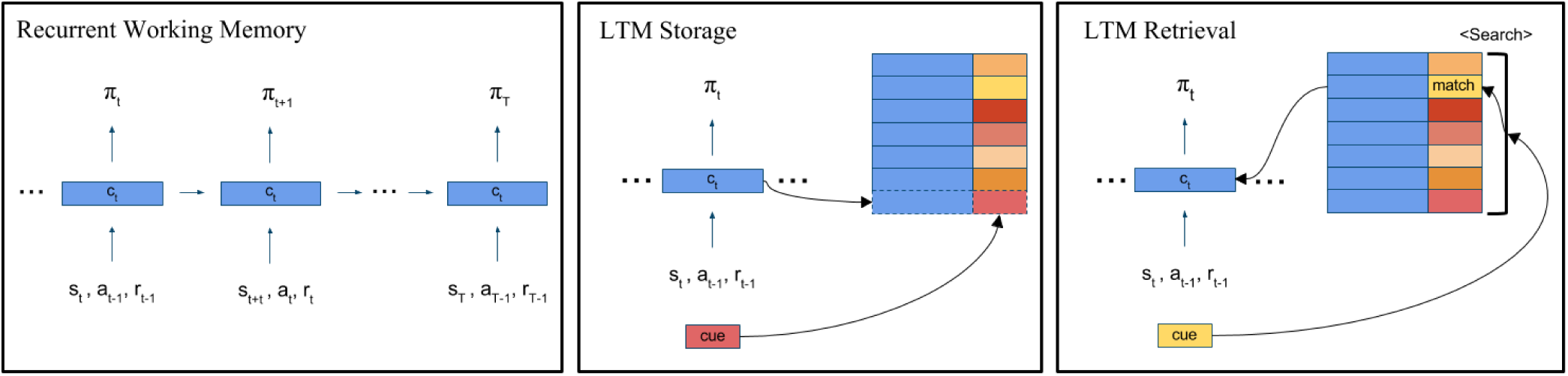
(Left) A high-level schematic of the recurrent network (LSTM) that comprises the episodic meta-RL (EMRL; Ritter et al., 2018) agent’s working memory. On each time step the LSTM receives an environment state, the action taken on the previous trial, and the reward received on the previous trial. The LSTM encodes this information incrementally into its cell state *c*, and then outputs a policy and value estimate (not shown). (Middle) The storage operation to long-term memory at a single LSTM time step. Storage is triggered when reward is received at the end of each two-step trial, at which point the agent appends a perceptual representation of the current context (a “cue”) along with its cell state to a non-parametric store of such items. (Right) The long-term memory retrieval operation which occurs on every time step. A search is carried out over the cues stored in long-term memory for the closest match to the current cue. The working memory activations associated with the closest match are retrieved and reinstated to the working memory state.

The retrieved activations are reinstated through a learned gating function that arbitrates among the influences on the current working memory state of (1) current perceptual inputs, (2) the previous working memory state, and (3) the working memory state retrieved from long-term memory. This gating mechanism is a natural extension to the standard LSTM working memory, which uses gates to arbitrate between new inputs and the previous working memory state:

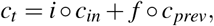

where *c_t_* is the current working memory state, *c_in_* is the agent’s representation of its current input, *c_prev_* is the working memory activation from the previous timestep, and o signifies elementwise multiplication.

The gates *i* and *f* are values between zero and one that allow (or disallow) inputs and and past working memory activations into the current state. These gates are computed accordingly:

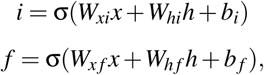

where *x* is the perceptual input, *h* is a function of the previous working memory state, and the weight matrices *W* and bias vectors *b* contain learned parameters.

To reinstate working memory activations retrieved from episodic memory without losing the current contents of working memory, our architecture adds the retrieved activations to the current working memory state, after passing them through a gate that is computed in a manner exactly analogous to the standard LSTM gates:

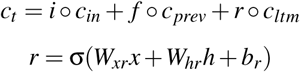

*c_ltm_* contains the retrieved activations from long-term memory. This reinstatement gate *r* is intended to learn to allow activations from episodic memory into working memory when they are useful, but not when they will interfere with the maintenance of important information in working memory.

To illustrate how this architecture works in practice, consider the episodic two-step task, wherein the working memory is expected to keep track of the reward probabilities at each outcome state. In order to infer these quantities, it must maintain information about recent actions and states. When the agent receives reward in the final step of a two-step trial, it will save the activations of its current working memory - which encode the agent’s outcome state and reward - to its long-term memory. These are saved along with a representation of the context stimulus. At the beginning of a future two-step trial, the agent will encounter this same stimulus. It will then search its long-term memory for matches for that stimulus, and will retrieve the hidden state from the past trial. Crucially, this hidden state will encode the state the agent encountered at the end of the last exposure to that stimulus as well as the reward received and the action taken. Possessing this critical episodic information, the agent is able to exploit the structure of the episodic two-step task. Specifically, it can learn to implement model-based or model-free valuation with respect to trial information retrieved from long-term memory.

## Methods and Results

After training, we assessed EMRL’s performance on a set of evaluation episodes which followed the same structure as the training episodes. To isolate the behavior of the *learned* learning algorithm operating in the activations, all data shown and described in this section were obtained with the weights frozen. To verify that the observed effects are consistent across trained agents, we performed the following analyses on 10 instances of each agent type, where each instance was trained with a different random seed. The reported statistical tests measure the variability across these agent instances. Agent optimization was performed by an implementation of asynchronous actor-critic (Mnih et al., 2016).

First, to determine whether EMRL’s episodic memory was performing effectively, we compared the reward EMRL acquired with that acquired by a control meta-RL agent (MRL; Figure 3a) that was trained and tested in exactly the same way, but did not have access to episodic memory (i.e., its r-gate was always fixed to zero). EMRL achieved more reward overall than MRL (t(9)=4.28, p=1.21e-4). This increase in reward came entirely from the cued trials: the difference between reward obtained by EMRL and that obtained by MRL in the cued trials was highly significant (t(9)=7.205, p=1e-6), while the difference in uncued trials was not significant (t(9)=-1.09, p=0.289). These results provide evidence that EMRL was able to use its episodic memory to acquire the additional reward available during cued trials.

**Figure 3:**
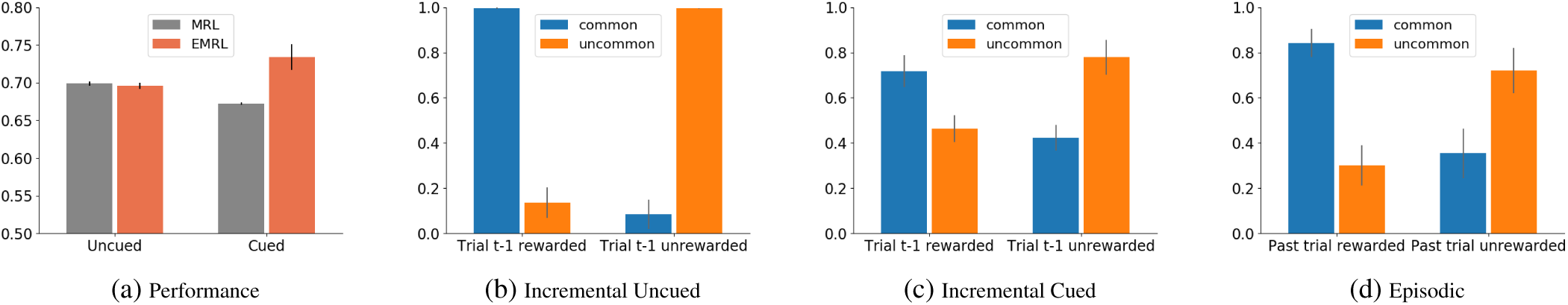
EMRL achieves more reward than MRL and exhibits both incremental and episodic model-based behavior. Bar heights indicate means over 10 trained agents with different random seeds. Error bars indicate the standard error of the mean over the 10 trained agents. (a) Average reward obtained by MRL and EMRL on cued and uncued trials. EMRL earns more reward than MRL on cued trials, suggesting that EMRL can use its episodic memory to exploit the task’s episodic structure. For comparison, a random policy achieves 0.5 reward on this task. (b) Proportion of uncued trials in which EMRL repeated the action it took on the previous trial (*t* − 1), split by whether it received reward on *t* − 1 and whether the transition on *t* − 1 was common. The interaction between those two factors is a sign of model-based learning^1^. (c) Same as b, but for cued trials. (d) Same as b, but split by whether EMRL received reward on the past trial *k* when the cue was first encountered and whether the transition on trial *k* was common.

Next, we asked whether EMRL exhibited the canonical patterns of incremental model-based and model-free behavior first described in the two-step task by (Daw et al., 2011). We formally tested for these patterns of behavior by performing ANOVAs on the probabilities of repeating the previous action, with two binary factors: whether the previous trial was rewarded, and whether the previous trial had a common transition. A main effect of previous trial being rewarded would indicate a model-free strategy, while an interaction between previous trial being rewarded and previous trial being common would indicate a model-based strategy^1^. On uncued trials (Figure 3b), we found a strong effect of the interaction term (F(1,9)=842, p=3.34e-10), indicating that the learned algorithm correctly exploited the transition structure of the task when no episodic information was available. This behavior reproduces the main two-step task result from Wang et al. (2017). The main effect of reward was not significant, indicating the absence of model-free behavior (F(1,9)=3.462, p=0.096). The main effect of transition type was also not significant F(1,9)=2.944, p=0.120). On cued trials (Figure 3c), we also found an effect of the interaction term (F(1,9)=53.78, p=4.4e-5), indicating that EMRL continued to use the incremental strategy during the cued trials. The main effect of reward was not significant, indicating the absence of model-free behavior (F(1,9)=0.170, p=0.690). The main effect of transition type was also not significant (F(1,9)=5.017, p=0.052).

Next, most centrally, we asked whether EMRL could apply model-based reasoning to information retrieved from episodic memory (Figure 3d). We performed the same ANOVA described above, but using as factors: whether the past cue-associated trial was rewarded and whether the past cue-associated trial had a common transition. Since our task guaranteed receiving the same reward if the agent reached the same state as the past trial, the agent should prefer to take the opposite action as on the past trial if it experienced an uncommon transition and received reward on that trial. We indeed found a strong effect of the interaction term in this analysis (F(1,9)=36.1, p=2e-4), and non-significant main effects of reward and transition type (F(1,9)=4.354, p=0.665; F(1,9)=4.132, p=0.073). We only performed this analysis on cued trials because the factors are undefined on uncued trials.

To supplement the ANOVA analysis, we fit a probabilistic choice model to EMRL’s behavior, similar to the model that Vikbladh et al. (2017) fit to their human data. This model casts participants’ choices as a softmax of the weighted sum of valuations from the four valuation strategies. Fitting this model yields estimates of the weight on each strategy (see Supplement for full details). In accordance with the ANOVA results, binomial sign tests on the estimated weights (Figure 4) revealed a significant contribution of incremental model-based learning (cued trials, p=0.002; uncued trials, p=0.002) and episodic model-based learning (cued trials, p=0.002), and no significant effect of incremental model-free learning (cued trials, p=0.754; uncued trials, p=0.109) or episodic model-free learning (cued trials, p=0.344).

**Figure 4:**
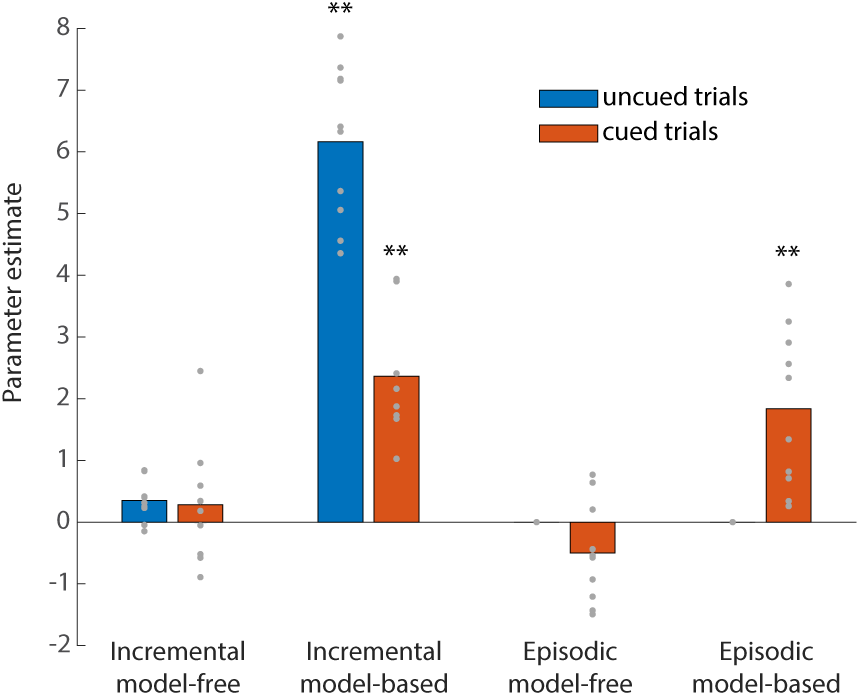
Parameter estimates in a model with weighted contributions of four decision systems. The estimates provide evidence that incremental and episodic model-based valuations contribute to EMRL’s behavior. The model was fit to EMRL’s actions separately on cued and uncued trials. Each gray dot denotes the model fit parameter for one of the 10 trained agents. ** denotes significant difference from 0 at p<0.005 by binomial sign test.

Overall, the ANOVA and model fit results confirm that EMRL exhibits both incremental and episodic model-based learning, but not episodic model-free learning, in accord with the human behavior observed by Vikbladh et al. (2017). The results show further that, like MRL (Wang et al., 2018), EMRL does not use incremental model-free learning, opting instead for the model-based incremental learning that achieves greater reward in the two-step task.

### Analysis of Reinstatement Gate Activations

To see what kind of gating function the r-gates learned, we carried out preliminary analyses of the r-gate activations. Figure 5a shows timecourses (averaged over 500 episodes) of the r-gate mean for a single trained agent. Each timecourse represents one two-step trial step; that is, whether it was a step where the agent took an action, saw an outcome, or received a reward. The timecourses show that the r-gate was more open during cued trials compared to uncued trials, consistent with the presence of useful episodic information on cued trials. Further, the r-gate was most open during the first stage of each trial. This is sensible, because this is the step on which reinstated information can be used to select an action. Figure 5b compares the r-gate mean for a single trained agent on cued trials when the agent selected the optimal or the opposite action. The r-gate was significantly more open on correct trials, consistent with the idea that retrieved information is necessary to select the correct action on these trials.

**Figure 5:**
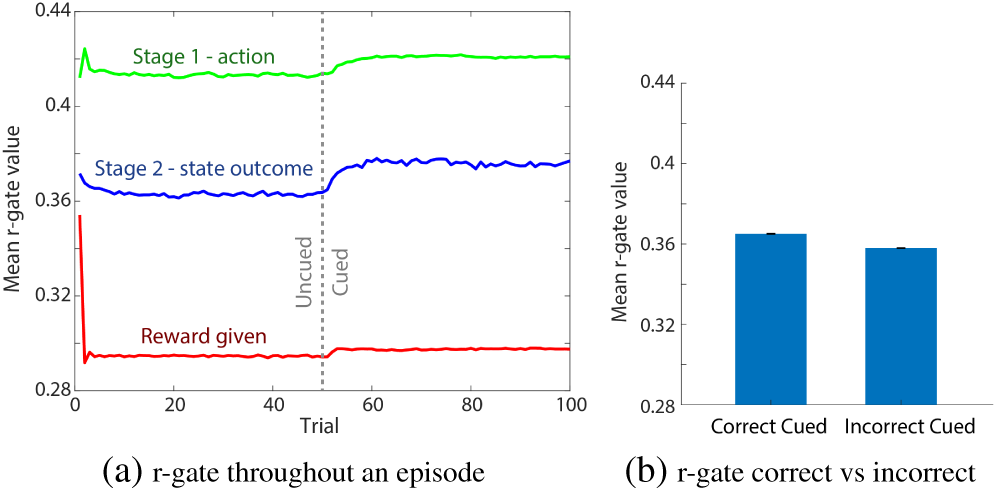
(a) Timecourse of the mean values of the reinstatement gate averaged over 500 episodes, split by stage of the trial. (b) Mean values of the reinstatement gate on cued trials on which the agent selected the optimal action (correct) and on cued trials on which it selected the opposite action (incorrect), averaged over all units.

While these effects are very clear, and the latter is highly significant, their magnitude is very small. Indeed, the r-gates allow a substantial proportion of the retrieved vector into working memory even during uncued trials when the information is not useful, and this proportion only increases slightly during the cued trials, where the information is critical. This suggests that mechanisms other than complete gating in/out are at play. For instance, the r-gates may subtly modify retrieved vectors to help the recurrent dynamics and policy layer determine whether or not to use the retrieved information. Further work will be needed to understand the solution the r-gates have found, and the insights gained may extend to biological gating systems. Alternatively, applying regularization, such as dropout, may lead to more easily interpretable neural mechanisms, as in Banino et al. (2018).

## Discussion

The experiments in this work establish that when trained via model-free learning on a task distribution with both incremental and episodic reward structure, EMRL learns to simultaneously execute incremental and episodic model-based learning algorithms. This ability to deploy and coordinate both learning algorithms mirrors that of the participants in the study by Vikbladh and colleagues, providing initial support for EMRL as a model of human decision making. EMRL thus provides an empirically grounded unified account of incremental and episodic learning processes, whereby a single model-free learning mechanism learns to execute and deploy the variety of learning algorithms observed in humans. In addition to support from behavioral data, the model accords in principle with a large neuroscientific literature supporting the notion that episodic memory retrieval recreates patterns of activity in neural circuits supporting working memory (Cohen & O’Reilly, 1996; Staresina, Henson, Kriegeskorte, & Alink, 2012; Hoskin et al., 2017; Xiao et al., 2017). Further, EMRL’s reinstatement gating mechanism fits naturally with well-supported theories input, output, and reallocation gating in prefrontal cortex (e. g., O’Reilly & Frank, 2006; Chatham & Badre, 2015).

It is worth pointing out *why* EMRL learns to use the valuation strategies that it does, and does not learn to use the others (summarized in Figure 4). In essence, optimized neural systems like MRL and EMRL will approach optimal behavior on their training distribution, subject to the limitations of the architecture design and efficacy of stochastic gradient learning. these experiments, the training distribution was the episodic two-step task itself, which yields greater reward to model-based than model-free strategies. EMRL’s architecture, with essential features such as associative retrieval and differentiable working memory, enables a broad spectrum of strategies – including model-based and model-free – to be learned via gradient descent. Accordingly, EMRL learns to use model-based instead of model-free strategies because (1) the architecture supports both strategies and (2) the model-based strategies earn more reward on the training distribution. Future work may investigate ecologically inspired training distributions (Anderson, 1990) that lead generic learning algorithms like MRL/EMRL to reproduce human-like deviations from optimality in laboratory tasks.

The key takeaway from the success so far of EMRL is a proof of the sufficiency of a small set of well motivated architectural components, when trained to optimize a specific objective function, to produce a variety of episodic and incremental learning processes observed in humans. The architecture components are: (1) a recurrent working memory with (2) a non-parametric store of working memory activations that can be retrieved by context and reinstated through (3) a learned gating system. The objective function is total reward achieved on a distribution of learning tasks which contain both incremental and episodic structure.

This model makes a number of predictions which may be tested through further empirical work:

- Pattern reinstatement has been observed during the retrieval of static stimuli (e.g., Bornstein & Norman, 2017); EMRL predicts that such pattern reinstatement should also be measurable during the retrieval of memories that encode sequences of events or the results of computation carried out in WM.
- There is considerable evidence that gating mechanisms regulate the flow of information into and out of working memory (Chatham & Badre, 2015). Analogous experiments should find evidence for a similar mechanism that gates reinstated activations from long-term memory into working memory.
- It should be possible to modulate the degree of WM pattern reinstatement on a given task by training participants on variants of that task where memory retrieval is either useful or distracting. EMRL predicts specifically that pattern reinstatement in PFC should be affected by such manipulations, while reinstatement in hippocampus should be unaffected.

In summary, this work presents a new theory that explains the collage of learning processes observed in humans during decision making as an interplay between working and episodic memory that is itself learned through training to maximize reward on a distribution of learning tasks. Future work may test the predictions made by this model and test the model’s ability to replicate additional sources of empirical data.

## Acknowledgements

We would like to thank Siddhant Jayakumar, Charles Blundell, and Razvan Pascanu for helpful discussion and advice; the CogSci reviewers for insightful comments and feedback; and those at Google and DeepMind behind the codebases and infrastructure we used to build the agents.

## Supplement: Model Fit Details

The probability of the agent taking action *a*_0_ in the starting state *s*_0_ at trial *t* was modeled as the softmax of the weighted sum of the differences between the value estimates for the two actions for each of the four valuation strategies:

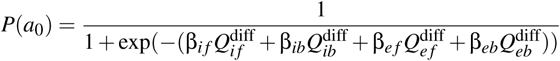

where *Q*^diff^ = *Q*(*s*_0_,*a*_0_) – *Q*(*s*_0_,*a*_1_). At each trial *t*, the incremental model-free value estimate *Q_if_* (*s*_0_,*a*) for the action *a* taken on trial *t* was updated as

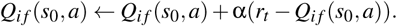

The incremental model-based value estimate *Q_ib_*(*s*_0_,*a*) was computed as

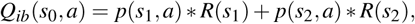

where *s*_1_ and *s*_2_ are the two second-stage states, *R*(*s*) is the agent’s estimate of the reward received in state *s* and *p*(*s,a*) is the probability of transitioning from state *s*_0_ into state *s* after taking action *a. R*(*s*) was updated as

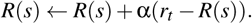

The episodic model-free value estimate *Q_ef_* (*s*_0_,*a*) was set equal to 1 if on the past (cue-associated) trial, the agent took action *a* and was rewarded, or took the other action and was not rewarded; *Q_ef_* (*s*_0_,*a*) was 0 otherwise. *Q_eb_*(*s*_0_,*a*) was 1 if on the past (cue-associated) trial, the agent took action *a* and this resulted in a common transition with reward or an uncommon transition without reward, or if the agent took the other action and this resulted in a common transition without reward or an uncommon transition with reward. *Q_eb_*(*s*_0_,*a*) was 0 otherwise.

All incrementally learned values were updated with the same learning rate α, for a total of five parameters. These were estimated by maximum likelihood on the concatenated data of all 500 episodes, with incrementally learned values reset to 0.5 at the beginning of each episode. Cued and uncued trials were fit separately. Note that episodic valuation is undefined during uncued trials.

## Footnotes

1 Akam, Costa, and Dayan (2015) demonstrated that there exist sophisticated model-free strategies which can, under the right conditions, produce this reward-by-transition interaction. See Wang et al. (2018), Supp. Figures 6 and 7 for an in depth analysis of this issue with respect to MRL.

## References

Akam, T., Costa, R., & Dayan, P. (2015). Simple plans or sophisticated habits? state, transition and learning interactions in the two-step task. PLoS Comput Biol, 11(12), e1004648.

Anderson, J. R. (1990). The adaptive character of thought. Psychology Press.

Banino, A., Barry, C., Uria, B., Blundell, C., Lillicrap, T., Mirowski, P.,…others (2018). Vector-based navigation using grid-like representations in artificial agents. Nature, 557(7705), 429.

Blundell, C., Uria, B., Pritzel, A., Li, Y., Ruderman, A., Leibo, J. Z.,…Hassabis, D. (2016). Model-free episodic control. *arXiv preprint* arXiv:1606.04460.

Bornstein, A. M., Khaw, M. W., Shohamy, D., & Daw, N. D. (2017). Reminders of past choices bias decisions for reward in humans. Nature Communications, 8.

Bornstein, A. M., & Norman, K. A. (2017). Reinstated episodic context guides sampling-based decisions for reward. Nature Neuroscience, 20.

Chatham, C. H., & Badre, D. (2015). Multiple gates on working memory. Current Opinion in Behavioral Sciences.

Cohen, J. D., & O’Reilly, R. C. (1996). A preliminary theory of the interactions between prefrontal cortex and hippocampus that contribute to planning and prospective memory. In Prospective memory: Theory and applications.

Daw, N. D., Gershman, S. J., Seymour, B., Dayan, P., & Dolan, R. J. (2011). Model-based influences on humans’ choices and striatal prediction errors. Neuron, 69(6), 1204–1215.

Dolan, R. J., & Dayan, P. (2013). Goals and habits in the brain. Neuron, 80(2), 312–325.

Duncan, K. D., & Shohamy, D. (2016). Memory states influence value-based decisions. Journal of Experimental Psychology: General, 145(11), 1420.

Gershman, S. J., & Daw, N. D. (2017). Reinforcement learning and episodic memory in humans and animals: An integrative framework. Annual review of psychology, 68.

Graves, A., Wayne, G., Reynolds, M., Harley, T., Danihelka, I., Grabska-Barwińska, A.,…others (2016). Hybrid computing using a neural network with dynamic external memory. Nature.

Hochreiter, S., & Schmidhuber, J. (1997). Long short-term memory. Neural computation, 9(8), 1735–1780.

Hoskin, A. N., Bornstein, A. M., Norman, K. A., & Cohen, J. D. (2017). Refresh my memory: Episodic memory reinstatements intrude on working memory maintenance. bioRxiv, 170720.

Lengyel, M., & Dayan, P. (2007). Hippocampal contributions to control: The third way. In Proc. of neural information processing systems, NIPS.

Mnih, V., Badia, A. P., Mirza, M., Graves, A., Lillicrap, T. P., Harley, T.,…Kavukcuoglu, K. (2016). Asynchronous methods for deep reinforcement learning. In Proc. of int’l conf. on machine learning, ICML.

O’Reilly, R. C., & Frank, M. J. (2006). Making working memory work: a computational model of learning in the prefrontal cortex and basal ganglia. Neural computation, 18(2), 283–328.

Pritzel, A., Uria, B., Srinivasan, S., Puigdomènech, A., Vinyals, O., Hassabis, D.,…Blundell, C. (2017). Neural episodic control. *arXiv preprint* arXiv:1703.01988.

Ritter, S., Wang, J. X., Kurth-Nelson, Z., Jayakumar, S. M., Blundell, C., Pascanu, R., & Botvinick, M. (2018). Been there, done that: Meta-learning with episodic recall. In Proceedings of the 35th international conference on machine learning.

Staresina, B. P., Henson, R. N., Kriegeskorte, N., & Alink, A. (2012). Episodic reinstatement in the medial temporal lobe. Journal of Neuroscience, 32(50), 18150–18156.

Sutton, R. S., & Barto, A. G. (1998). Reinforcement learning: An introduction (Vol. 1). MIT press Cambridge.

Vikbladh, O., Shohamy, D., & Daw, N. (2017). Episodic contributions to model-based reinforcement learning. In Annual conference on cognitive computational neuroscience, CCN.

Wang, J. X., Kurth-Nelson, Z., Kumaran, D., Tirumala, D., Soyer, H., Leibo, J. Z.,…Botvinick, M. (2018). Prefrontal cortex as a meta-reinforcement learning system. Nature neuroscience, 21(6), 860.

Wang, J. X., Kurth-Nelson, Z., Tirumala, D., Soyer, H., Leibo, J. Z., Munos, R.,…Botvinick, M. (2017). Learning to reinforcement learn. Retrieved from `arXivpreprint` arXiv: 1611.05763

Wimmer, G. E., Braun, E. K., Daw, N. D., & Shohamy, D. (2014). Episodic memory encoding interferes with reward learning and decreases striatal prediction errors. J Neurosci.

Wimmer, G. E., & Buechel, C. (2016). Reactivation of reward-related patterns from single past episodes supports memory-based decision making. Journal of Neuroscience, 36(10), 2868–2880.

Xiao, X., Dong, Q., Gao, J., Men, W., Poldrack, R. A., & Xue, G. (2017). Transformed neural pattern reinstatement during episodic memory retrieval. Journal of Neuroscience, 37.

